# Selective Modulation of Heart and Respiration by Optical Control of Vagus Nerve Axons Innervating the Heart

**DOI:** 10.1101/2020.06.26.173898

**Authors:** Arjun K. Fontaine, Gregory L. Futia, Pradeep S. Rajendran, Samuel Littich, Naoko Mizoguchi, Kalyanam Shivkumar, Jeffrey L. Ardell, Diego Restrepo, John H. Caldwell, Emily A. Gibson, Richard F. Weir

**Author notes:** Co-first authors (Arjun K. Fontaine, Gregory L. Futia). Co-senior authors (Diego Restrepo, John H. Caldwell, Emily A. Gibson, Richard F. Weir).

## Abstract

Targeting specifics subsets of peripheral pathways of the autonomic nervous system will enable new avenues to study organ control and develop new disease therapies. Vagus nerve stimulation (VNS) has shown many therapeutic benefits but current approaches involve imprecise electrical stimulation that gives rise to adverse effects, and the functionally relevant pathways are poorly understood. One method to overcome these limitations is the use of optogenetic techniques, which facilitate highly specific neural communication with light-sensitive actuators (opsins). Opsins can be targeted to cell populations of interest based on the location of viral delivery and genetic control of expression. Here, we tested whether holographic photostimulation of subsets of axons of the cervical vagus nerve that innervate the heart can be used to modulate cardiac function. Viral injection of retrograde adeno-associated virus (rAAV2-retro) in the heart resulted in robust, primarily afferent, opsin reporter expression in the vagus nerve, nodose ganglion, and brainstem. Selective holographic photostimulation of axons resulted in changes in heart rate, surface cardiac electrogram, and respiratory responses that were different from responses elicited by whole nerve photostimulation.

---

Improved techniques are necessary for selective modulation of peripheral nerve fibers, particularly within the autonomic nervous system. Effort in recent years has increasingly been aimed at targeting neural pathways of abdominal and thoracic organs for therapeutic neuromodulation^1–3^. While pharmacologic treatments have poorly confined spatial and temporal precision, the premise underlying initiatives in ‘bioelectronic medicine’ or ‘electroceuticals’ is compelling; tapping into the neural circuitry driving organ function may enable approaches for disease treatment pathways unachievable or far less precisely achievable with pharmacologic approaches.

Vagus nerve stimulation (VNS) interventions using electrode-based techniques have shown clinical efficacy in treating numerous diseases. VNS is clinically-approved for the treatment of epileptic seizures^4,5^ and depression^6^, and clinical studies demonstrate benefit in treating inflammatory disorders such as Crohn’s disease^7^ and rheumatoid arthritis^8^, as well as hypertension^9^ and obesity^10^. For the heart, clinical studies point to therapeutic potential in treating atrial arrhythmias^11,12^ and heart failure^13–15^, while preclinical data suggests vagal stimulation has cardioprotective effects in models of pressure and ischemia by mitigating intrinsic and sensory neuron remodeling and cardiac hypertrophy^16–18^.

However, the fiber types and mechanisms that underlie the functional effects of VNS are either unknown or poorly understood. Given the range of organs which are innervated by the vagus nerve and the non-selectivity of electrode stimulation, many off-target fibers are stimulated for the therapies mentioned above. Adverse effects arising from VNS therapy include pain, paresthesia, hoarseness, voice alteration, cough, dyspnea, nausea and headache^5,19,20^.

The limited mechanistic understanding and imprecise stimulation underlying current peripheral nerve therapies highlight the need for tools with greater selectivity and specificity. A potential avenue for achieving these improvements is optically mediated intervention using optogenetic tools^21^. With such techniques, light-driven activation or suppression of nerve pathways is possible with cell-type specific targeting^22^. In preclinical animal studies, virally-targeted excitatory opsins were employed in peripheral nerves to optically stimulate selective motor activity^23,24^, while inhibitory opsins have been used to suppress muscle activity^23,25^ and inhibit pain^26^. Adeno-associated viruses (AAVs) are widely used to deliver optical actuators and sensors in experimental models, with targeting specificity defined by tissue/location of delivery, virus serotype, and genetic promoter. There are numerous AAV-based gene therapies in human clinical trials, and to date, three have been approved for treatment in the U.S. and E.U.^27–30^.

Given the advantages in precision that optical techniques offer, and the encouraging progress of AAV gene delivery in the clinical setting, we have investigated the possibility of using retrograde AAV injection into an organ of interest to modulate its function with targeted optical stimulation in proximal nerve pathways. The present study focuses on the heart, which is regulated by extensive and complex control from the autonomic nervous system^31,32^. We demonstrate this approach with injection of a retrograde AAV into the heart to transduce opsins proximally in fibers of the vagus nerve. Selective photostimulation of subsets of heart-specific axons was applied at the cervical vagus nerve using both one-photon illumination and two-photon holographic excitation^33,34^. Significant changes in heart rate and electrocardiogram (ECG) parameters, as well as the elicitation of respiratory reflex, serve as a proof-of-concept for organ-specific optogenetic neuromodulation and study of organ function.

## Methods

### Heart Surgery and Viral Injection

The use of animals was approved by the Institutional Animal Care and Use Committee (IACUC) at the University of Colorado, Anschutz Medical Campus, with accreditation by the Association for Assessment and Accreditation of Laboratory Animal Care (AAALAC). All experiments were performed in accordance with IACUC regulations under an approved protocol. *pAAV-Syn-Chronos-GFP* (59170-AAVrg)^35^, *pAAV-Syn-ChR2(H134R)-GFP* (58880-AAVrg)^36^, and *AAV-CAG-hChR2-H134R-tdTomato* (28017-AAVrg)^37^ were packaged in AAV-retrograde (rAAV2-retro), a virus designed for efficient retrograde access to projection neurons^38^, and obtained from Addgene. The injected titers were approximately ∼1×10^12 vg/mL, and a total of 20µl volume was injected in each animal/procedure.

Mice (C57BL/6) were given carprofen (5 mg kg^-1^, s.c.) and buprenorphine (0.05 mg kg^-1^, s.c.) 1 hour prior to surgery. Animals were anesthetized with isoflurane (induction at 5%, maintenance at 1-3%, inhalation), intubated, and mechanically ventilated (CWE Inc., SAR-830/AP). Core body temperature was measured and maintained at 37°C. The surgical incision site was cleaned 3 times with 10% povidone iodine and 70% ethanol in H_2_O (vol/vol). A left lateral thoracotomy was performed at the fourth intercostal space, the pericardium opened, and the heart was exposed. One microliter AAV stock (∼1×10^13 vg/mL) diluted in 10 uL 0.01 M phosphate-buffered saline was subepicardially injected into *each* the left and right ventricle with a 31-gauge needle. The surgical wounds were closed with 6-0 sutures. Buprenorphine (0.05 mg kg^-1^, s.c.) was administered once daily for up to 2 days after surgery.

### Vitals Measurement

Mouse vitals (heart rate, pulse distension, breath rate, breath distension, pO2) were monitored and recorded in anesthetized mice with a MouseOx Plus suite (Starr Life Sciences) with a paw sensor. For ECG measurement, three needle electrodes were placed subdermally: one on both sides of the upper chest cavity near the armpits and lead I in the lower abdomen. Signals were amplified (BioAmp) and digitized using a PowerLab 4/35 and were analyzed in LabChart software (ADInstruments).

#### 1 Photon Photostimulation of Cervical Vagus Nerve

Mice were tested in terminal procedures over a period of 6-23 weeks post viral injection, with robust expression and functional responses observed at 23 weeks. Animals were anesthetized using 1-3% isoflurane gas, maintained through nose cone inhalation, and placed supine on a heating pad, while vitals were monitored throughout the experiment. A 1 cm incision was made at the cervical region, 2-3 mm lateral to midline on the mouse’s left side. Blunt dissection techniques were used to expose the left cervical vagus nerve and separate it from the carotid artery. An optical cannula (CFMC12L20, Thorlabs) was positioned with a micromanipulator to abut the nerve for laser photostimulation. A 473nm-wavelength solid-state laser (BL473T8-150FC, Shanghai Laser & Optics Century) was coupled to the optical cannula with an optical patch cable (200µm core, .39 NA, M81L005, Thorlabs) and interconnect (ADAF2, Thorlabs). The laser was controlled with a custom Arduino circuit board to output 5ms pulses at 20 Hz.

### GRIN Lens-Integrated Cervical Nerve Cuff

The nerve cuff was fabricated from silicone (Silastic MDX4-4210, Dow Corning) by compression molding. Mold components were designed in SolidWorks and fabricated using Direct Laser Metal Sintering (DLMS) in maraging steel. A GRIN lens, 9 mm in length, 1 mm in diameter (Inscopix 1050-002177), was placed in a polyimide tube (7.5 mm length, 1 mm inner diameter, 1.15mm outer diameter, Microlumen Inc.) and fixed with cyanoacrylate. This assembly was inserted into the silicone cuff at a perpendicular orientation to the nerve’s longitudinal axis, with the GRIN lens inset 250 µm from the interior surface of the cuff, and fixed with cyanoacrylate. The cuff was fully assembled before implanting in the animal (Figure1A,B).

**Figure 1:**
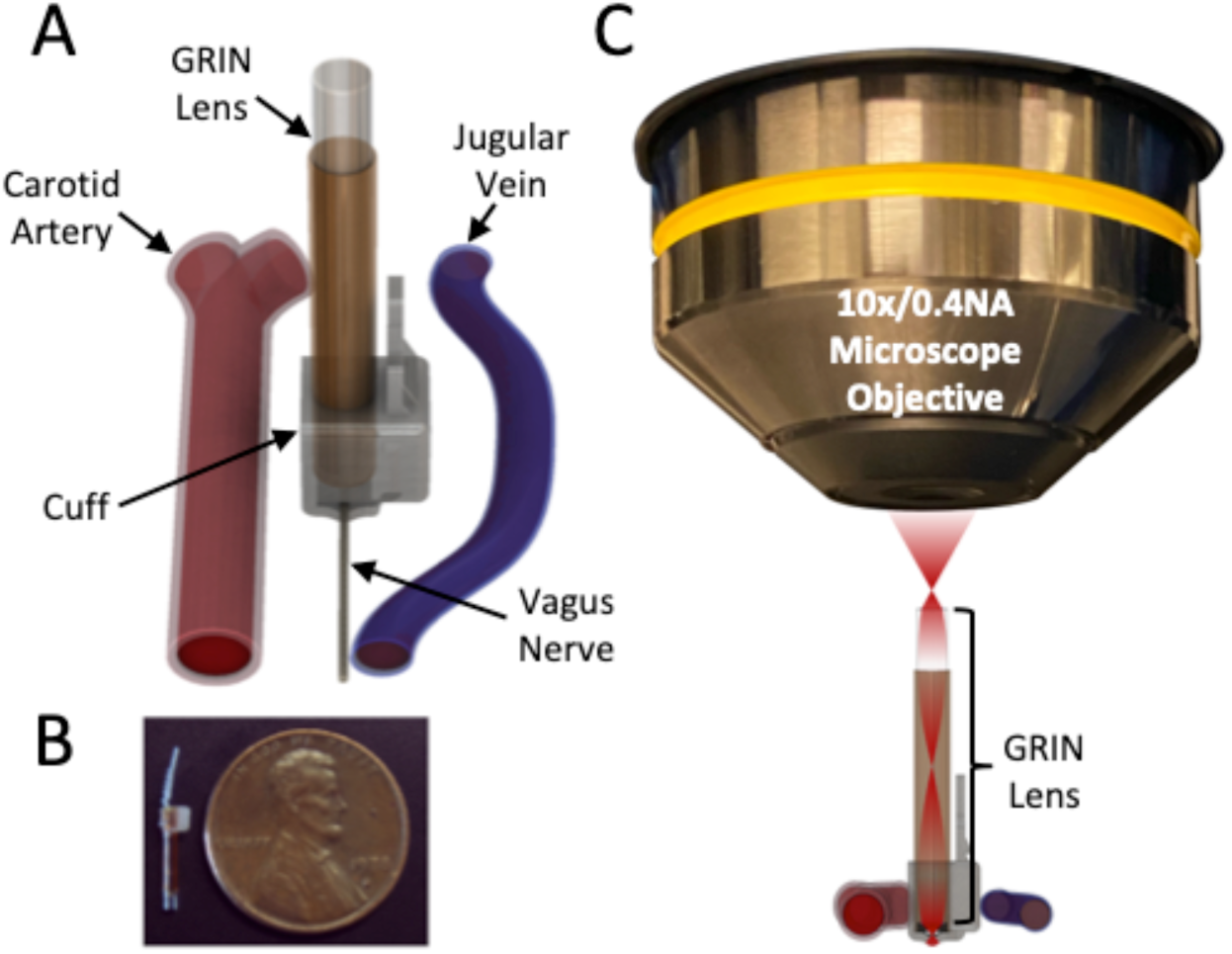
Detailed schematic of the GRIN-lens-integrated nerve cuff for two-photon imaging and holographic stimulation. (A) GRIN-cuff situated on the cervical vagus nerve between the carotid artery and jugular vein. (B) Photo of device next to penny for scale, in the un-cuffed (non-ratcheted) position (C) Schematic of GRIN-cuff and microscope objective configuration.

#### 2 Photon Imaging and Holographic Photostimulation

The GRIN lens-integrated nerve cuff was fastened to the vagus nerve of the anesthetized mouse with a ratcheting strap and held rigid using custom-modified dental forceps mounted to a three-axis manipulator. The mouse was placed supine with the GRIN lens positioned under the objective lens of an upright microscope for two photon imaging and holographic photostimulation (Figure1C).

A custom setup for imaging and holographic photostimulation was built on a Movable Objective Microscope (Sutter Instruments, Inc.), as described previously^34^ and shown schematically in Supplementary Figure1. A Ti:Sapphire laser oscillator (MaiTai HP, Newport, Inc.) producing ∼80 fs pulses at 920 nm with an 80 MHz repetition rate provided the excitation light for two-photon imaging. The GRIN lens (0.5 NA, 1:1 magnification) served as an optical relay, such that the cervical vagus nerve was in the conjugate plane at the focus of the objective lens (0.4 NA, Olympus USLSAPO10×2). Fluorescence emission from the sample was collected in the epi-direction, filtered by a dichroic (LP665) and bandpass filter (green channel, 510/42 and red channel, 620/60) and detected on large-area photomultipliers (10770PA-40, Hamamatsu). Typical imaging frame rates were 3.5-5.5 Hz with an imaging field of view of ∼ 200 μm.

Two-photon photostimulation was performed using a 1030 nm pulsed laser (SpectraPhysics, Spirit HE 1030-70) producing pulses with ∼240 fs pulse duration at 500 kHz repetition rate. The photostimulation beam path included a spatial light modulator (SLM, Phasor, 3i, Inc.) to create a desired spatial pattern at the objective focus^39^. The 1030 nm laser beam was expanded to fill the active area of the SLM and adjustable phase patterns were calculated in software (Slidebook 6, 3i, Inc.) and output onto the SLM. The 1030 nm laser was combined with the 920 nm laser using a polarization beam splitter placed directly after the scan lens. The SLM was imaged onto the back aperture of the objective lens at 1.6X magnification to slightly over fill the aperture. The phase pattern imparted on the beam by the SLM results in an intensity pattern at the focus of the objective. The spatial pattern at the sample was verified using a flat fluorescence slide detected on a widefield imaging system. An integrated acousto-optic modulator (AOM) in the 1030 nm laser was triggered with a function generator to control the timing for photostimulation. The setup is optimized for two-photon photostimulation of red-shifted opsins such as Chronos or ChrimsonR combined with imaging of reporter proteins such as GCaMP6. Additionally, the same setup was utilized for photostimulation at 920 nm, for channelrhodopsin (ChR2). Here, the signal output from a noncollinear optical parametric amplifier (NOPA-VIS-IR, Newport, Inc) tuned to 920 nm with a pulse duration of ∼50 fs and repetition rate of 500 kHz, was sent through the same SLM beam path for holographic photostimulation.

### GRIN Lens Aberrations

The GRIN relay lens introduces aberrations that can distort the laser focus off-axis. For imaging, this results in a fall off in intensity off axis. The typical field of view observed was approximately 200 μm in diameter, encompassing the full mouse cervical vagus nerve. Aberrations also expand the axial confinement of the holographic stimulation and this becomes worse for photostimulation regions which are off axis^40^. We fixed the spacing from the imaging plane in the nerve to the GRIN lens surface at the optimal working distance of 300 μm using the cuff. However, any deviation from the working distance can introduce further aberrations and reduce axial confinement^41^.

### Spatial Localization of 2-Photon Photostimulation

The two-photon excitation spatial profiles used for the animal experiments were characterized experimentally. A thin fluorescent sample on a slide was translated axially through the focus of the objective and GRIN relay lens and images of the fluorescent profile were acquired in a transmission microscope using a 10X 0.4 NA objective and imaged on a CMOS camera. In this manner, the spatial profile in x,y, and z were fully mapped. Supplementary Figure2 shows the profiles used for the two-photon photostimulation reported in the Results section. Note that the axial extent of excitation increases with increasing lateral illumination area when performing holographic stimulation^42^. Better axial sectioning can be accomplished using temporal focusing but was not used in these experiments^39^.

Scattering of light in biological tissue also plays a role in the intensity pattern in the nerve and can limit the achievable depth. Additionally, distortions in the wavefront due to the heterogeneous tissue will also change the spatial excitation pattern in the nerve. In the future, implementing a method to characterize the intensity *in vivo* such as the technique by Lerman *et al.*^43^ will be important for precise control of illumination in this type of application.

### Tissue Histology

Animals were euthanized with urethane (2 g/kg, i.p.). With a small-gauge needle inserted into the apex of the left ventricle, and incision made in the vena cava, animals were transcardially perfused with 50 ml ice-cold phosphate-buffered saline (PBS) containing 100U heparin followed by 50 ml of 4% paraformaldehyde (PFA) in PBS. Vagus nerves, nodose ganglia and superior cervical ganglia were excised and post-fixed in 4% PFA for 20 minutes, then rinsed in PBS. Tissues were slide-mounted with standard mounting media and imaged for examination of reporter expression on a confocal spinning disk microscope (Marianas, Intelligent Imaging Innovations).

Brains were harvested and post-fixed overnight on ice before cryoprotection by incubation in 0.1M PBS with 30% sucrose overnight at 4°C. Brains were placed in a positional mold and cut transversally (coronal sections) in a Leica cryostat (Leica CM1900, Leica Biosystems, Nussloch gmbh, Germany) at 30 um thickness. All sections were imaged on a Nikon A1R microscope (Tokyo, Japan) with a 40X objective 1.3 N.A.

Brain sections were washed three times in PBS before antigen retrieval permeabilization (sodium citrate 10 mM, Tween 0.5%, antigen retrieval pH 6.0, Bio wave 550W, 32°C, 5min). Sections were then permeabilized with 1% Triton X-100 in PBS for 30 min. Sections were washed three times in PBS between each permeabilization process. After permeabilization, all slices were subjected to a blocking step for 1hr in 5% normal donkey serum (NDS) including 0.3% tween in PBS before primary antibody incubation overnight at room temperature. The following primary antibodies were used and diluted in 5% NDS: goat anti-ChAT antibody (AB144P, 1:200) and chicken anti-GFP (Aves, GFP-1020, 1:500). Sections were washed three times in PBS before incubation in secondary antibodies diluted in 5% NDS for 2hrs at room temperature. The following secondary antibodies were used: donkey anti-goat Alexa Fluor 594 (1:500) and donkey anti-chicken Alexa Flor 488 (1:500). Sections were counterstained with Nissl (NeuroTrace™ 640/660 Deep-Red Fluorescent Nissl Stain - Solution in DMSO, 1:400) for 1hr at room temperature, and washed three times in PBS before sealing with a cover glass.

For whole-mount heart staining, fixed hearts were blocked in 0.01 M PBS with 10% NDS and 0.2% Triton X-100 PBS for 6 h at room temperature with agitation. Tissues were then incubated in primary antibody diluted in 0.01 M PBS with 0.2% Triton X-100 and 0.01% sodium azide for 3 nights at room temperature with agitation. The following primary antibody was used: chicken anti-GFP (Aves, GFP-1020, 1:1000). Tissues were washed several times in 0.01 M PBS overnight before incubation in secondary antibodies diluted in 0.01 M PBS with 0.2% Triton X-100 and 0.01% sodium azide for 2 nights at room temperature with agitation. The following secondary antibodies were used: donkey anti-rabbit Cy3 (Jackson ImmunoResearch, 711-165-152, 1:400), donkey anti-chicken 647 (Jackson ImmunoResearch, 703-605-155, 1:400), donkey anti-sheep Cy3 (Jackson ImmunoResearch, 713-165-003, 1:400), and donkey anti-goat Cy3 (Jackson ImmunoResearch, 705-165-003, 1:400). Whole hearts were washed several times in 0.01 M PBS overnight before being mounted on microscope slides in RIMS.

### Statistics

For statistical group comparisons a two-tailed two-sample unpaired t-test was used.

## Results

### One-Photon Cervical Photostimulation in ChAT-ChR2 Transgenic Mice

Heart rate and ECG parameters were analyzed in response to vagal photostimulation in ChAT-ChR2 transgenic mice (expressing the ChR2 opsin in cholinergic, and thus parasympathetic axons). With 473nm illumination at the left cervical vagus nerve, as expected^44,45^, heart rate was reduced substantially from baseline during stimulation and returned to baseline following the stimulus (−54%, p=8.1e-37)(Figure 2A). The Q-T interval also decreased significantly (−43%, p=8.0e-26), along with P-wave amplitude (−26%, p=4.3e-9) and T-wave amplitude (−13%, p=2.1e-13), while the P-R interval increased during stimulation (+5%, p=9.2e-12)(Figure2A&B).

**Figure 2:**
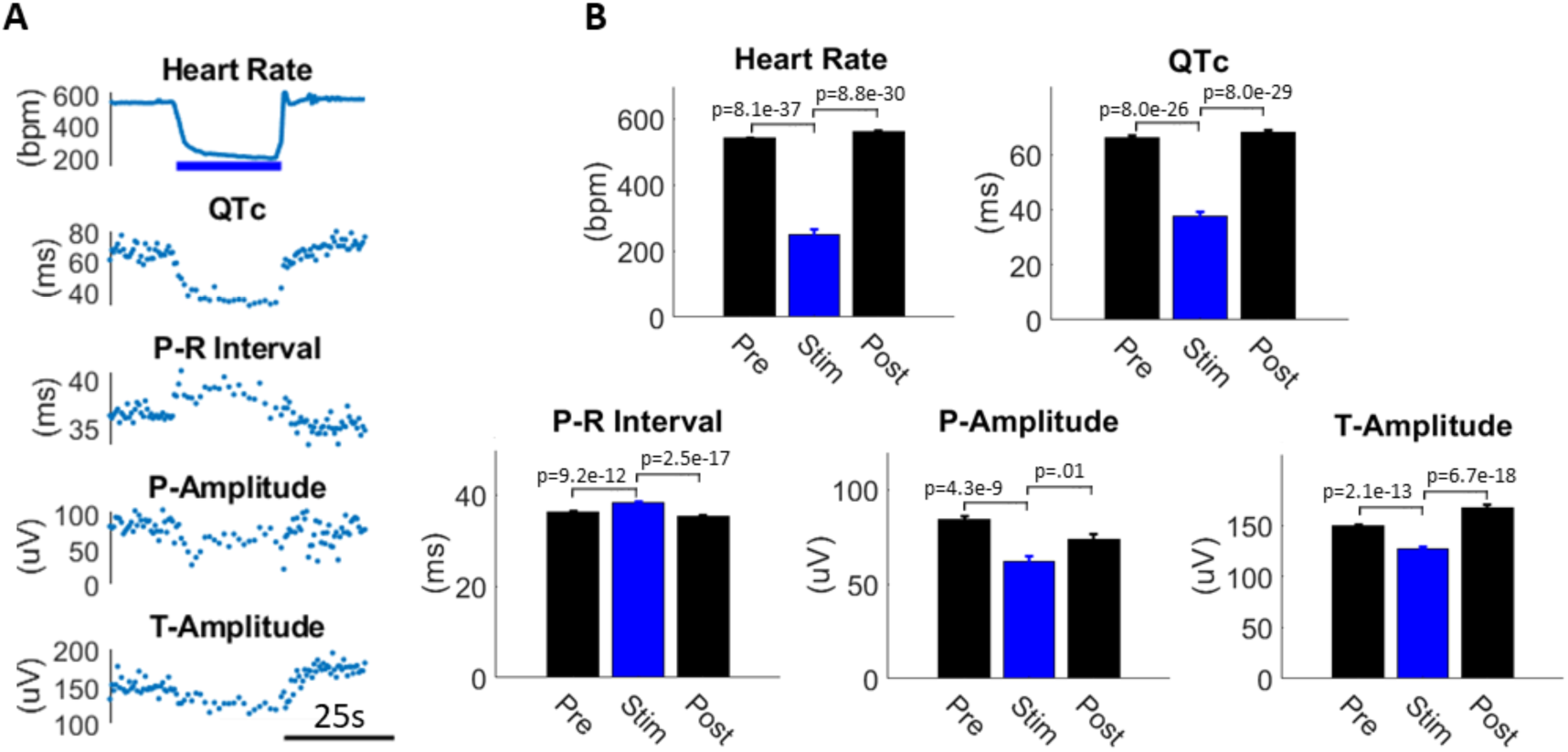
1-photon photostimulation of cervical vagus nerve in a ChAT-ChR2 transgenic mouse. (A) Heart rate is reduced, along with Q-T interval (QTc), P-wave amplitude and T-wave amplitude, while P-R interval is increased during stimulation. (B) Quantification of ECG parameters during the pre-stimulus period, stimulus period and post-stimulus period. (Blue bar: 473nm, 5ms pulses, 20Hz, 15mW unmodulated power)

### Retrograde Transduction of Nerve and Ganglia with Heart-Injected rAAV2-retro

Expression of the virally transduced opsin-fluorophore construct was observed in proximal nerve/ganglia targets for the three rAAV2-retro constructs injected: 1.) Syn-Chronos-eGFP, 2.) Syn-ChR2(h134R)-eGFP, and 3.) CAG-ChR2(h134R)-tdTomato (Figure3). A subset of axons in the left and right cervical vagi had robust reporter expression, and labeled axons appeared to be non-fasciculated and distributed across the nerve. A sparse subset of neurons were also labeled within the nodose ganglia, indicating the uptake in afferent (sensory) fibers. Superior cervical ganglia (SCG) were examined in a subset of mice, and all contained sparse neuronal labeling (Figure3B). Viral injections of construct #1 (above) were delivered in transgenic mice expressing ChAT-tdTomato to provide a cholinergic label, while the other constructs were injected in wild-type mice. In the ChAT-tdTomato-injected mice, viral expression was distributed across primarily non-cholinergic axons as well as in some ChAT-positive axons, suggesting an afferent-dominated expression profile (Figure3A). In these mice, vagi that were counted had an average of 25.2 ± 2.3 axons labeled with the viral reporter, of which 6.8 ± 1.7 (27%, n=3) were co-labeled with the transgenic ChAT reporter.

**Figure 3:**
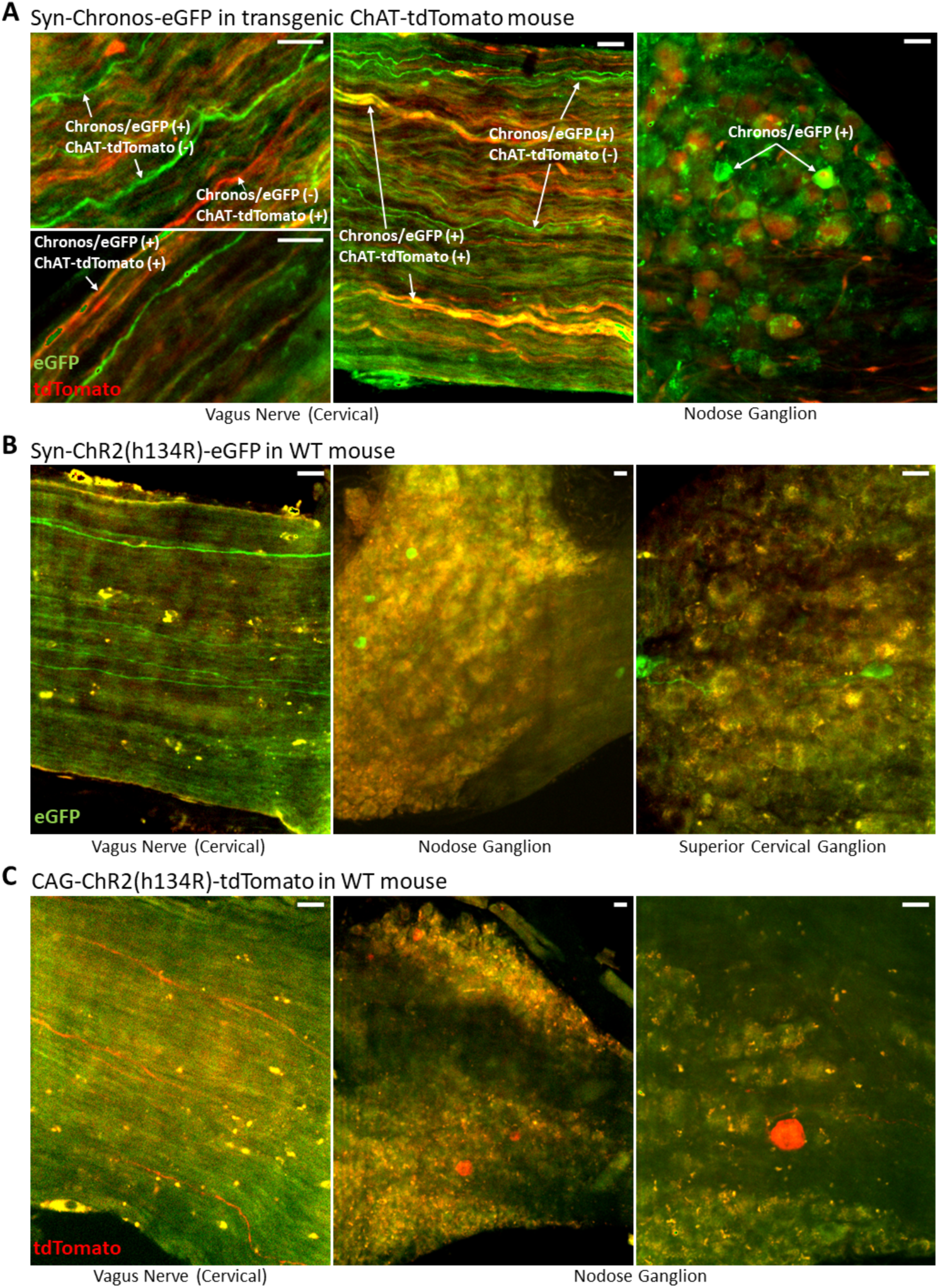
Retrograde labeling in nerve and ganglia with heart-injected rAAV2-retro. Three constructs were injected: (A) Syn-Chronos-eGFP, injected in ChAT-tdTomato transgenic mice. eGFP expression (green) was observed in ChAT-positive (tdTomato, red) and ChAT-negative axons throughout the vagus nerve, and in neurons of the nodose ganglion. (B) Syn-ChR2(h134R)-eGFP (green). eGFP expression is found in axons of the cervical vagus and nodose ganglia neurons, as well as in neurons of the SCG (C) CAG-ChR2(h134R)-tdTomato. tdTomato reporter expression (red) is observed in cervical vagus nerve and nodose ganglia, in a similar expression pattern. (all scale bars are 30µm)

### Retrograde Transduction in the Brainstem with Heart-Injected rAAV2-retro

rAAV2-retro reporter expression was observed in axons and a small number of cells in the nucleus of the solitary tract (NTS) of the dorsal brainstem (Figure 4) where we visualized cholinergic neurons by either expression of the transgenic ChAT-tdTomato reporter or by ChAT immunofluorescence with AlexaFluor594. Viral eGFP expression was found in axons, without cholinergic co-labeling, affirming the likelihood that these processes were afferent sensory projections (Figure 4A). Interestingly, some sparse cholinergic neurons in NTS were also labeled with eGFP (Figure 4B). Clear neuronal expression of viral reporters were not observed in the dorsal motor nucleus of the vagus (DMV, Figure 4A) or the nucleus ambiguous (NA, not shown). Reporter expression was also observed in the heart at the site of viral injection (Figure 4C).

**Figure 4:**
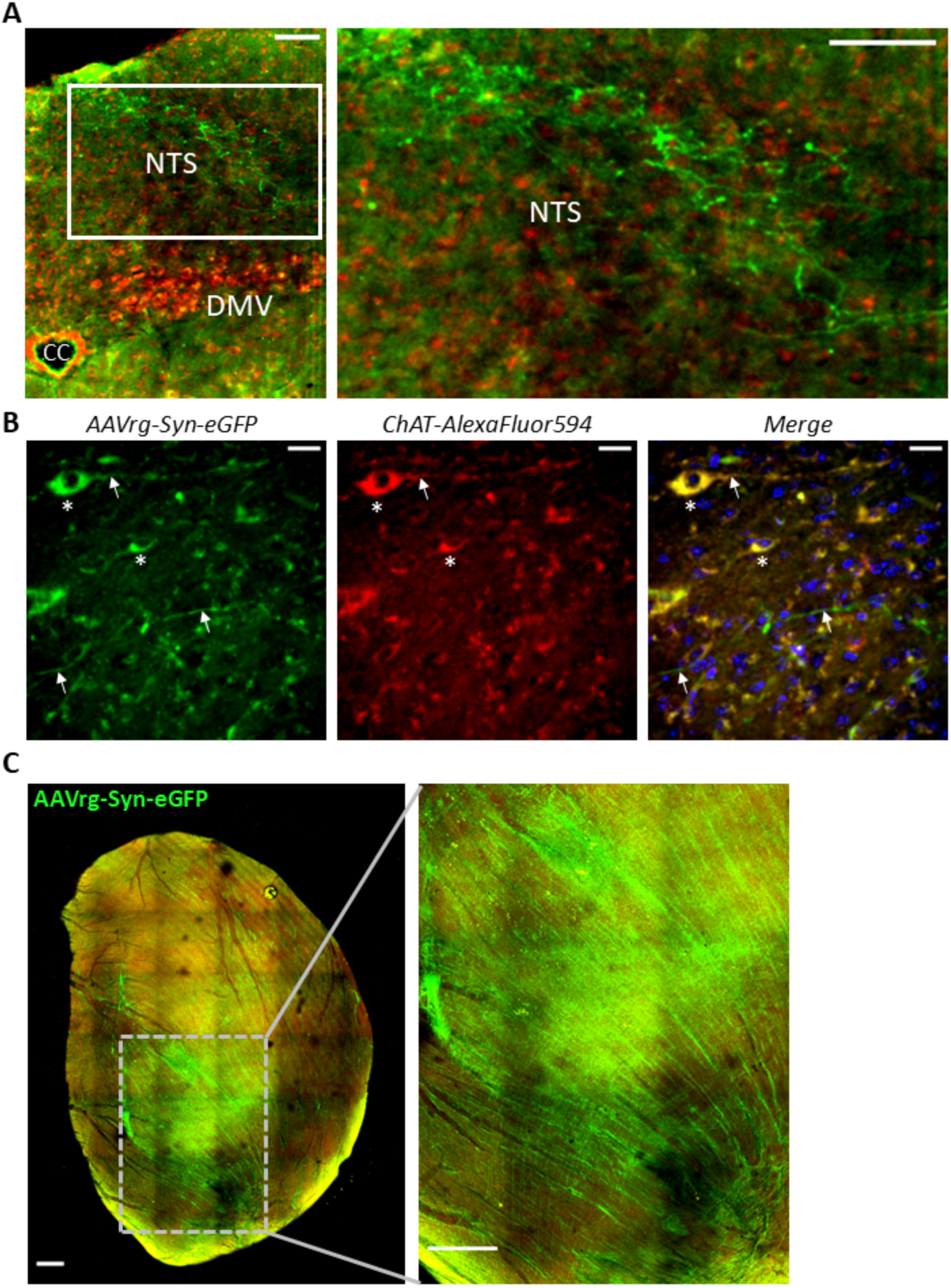
Retrograde viral reporter labeling in the dorsal brainstem and heart. (A) Representative confocal image showing eGFP in axonal processes (green) in the NTS in mice infected with AAVrg-Syn-ChR2(h134R)-eGFP. Cholinergic neurons of the dorsal motor nucleus of the vagus (DMV) were immunolabeled with ChAT-AlexaFluor594 (red). CC: central canal (scale bar, 100 µm) (B) Virally transduced neurons expressing eGFP within NTS were also observed. *Left:* At least two neurons (white asterisks) as well as axons (arrows) are positively labeled with eGFP. *Middle*: ChAT antibody stains the eGFP-positive neurons. *Right*: Merged image along with Hoechst stain (blue) shows the co-labeled cholinergic neurons (yellow) and non-co-labeled axons (green) (scale bar, 20 µm). (C) eGFP reporter expression (green) in a heart injected with AAVrg-Syn-Chronos-eGFP. Whole heart shown (*left*) with inset shown (*right*).

### Photostimulation of the Cervical Vagus Nerve in rAAV2-retro Heart-Injected Mice

One-photon photostimulation of the left cervical vagus nerve in anesthetized mice was performed with a 473nm laser via an optical fiber/cannula (Figure 5A). Here, stimulation was non-selective and activated axons throughout the nerve cross-section. Mouse vitals (heart rate, breath rate, breath distension) as well as ECG parameters, were perturbed by optical stimulation. Response profiles were variable across mice however, with both increases and decreases in heart rate observed, with or without respiratory changes. This variability in functional responses could be explained by inconsistent viral uptake across fiber types due to injection-to-injection site variability. An example response profile is shown in Figure 5; three consecutive stimuli ranging from 30-65 s in duration elicited reproducible, highly significant increases in heart rate and breath distension which arose during the optical stimulus and decayed to baseline 75-105 s following each stimulus (Figure 5B). The onset of breath distension increase was delayed 20-30 s relative to the onset of heart rate change. Significant changes in ECG waveform parameters (representative ECG waveform shown in Figure 5C) were observed concomitant with heart rate excursion (Figure 5D). ECG parameters were more sporadic during the perturbed heart rate, with mean levels changing significantly (Supplemental Figure1).

**Figure 5:**
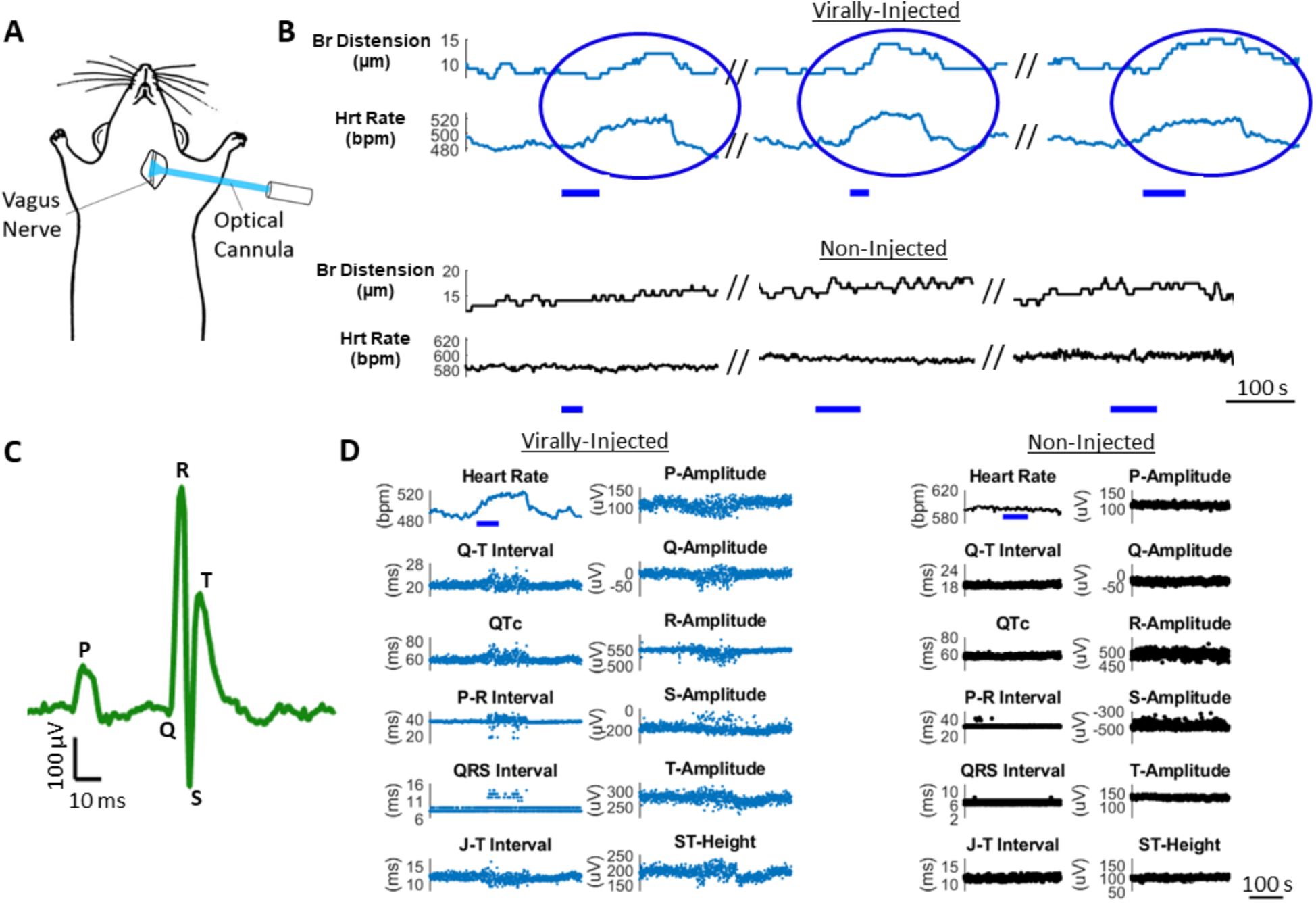
1-Photon stimulation of the cervical vagus nerve perturbs vitals and ECG parameters. (A) Schematic of photostimulation in the anesthetized mouse. The left vagus nerve is exposed and illuminated with a 473nm laser, delivered through an optical fiber/cannula. (B) Three consecutive photostimuli (blue bar) cause increases in heart rate and breath distension in a virally-injected (AAVrg-Syn-Chronos-eGFP) mouse while no changes occur in the non-injected control mouse. Note that the vertical axes in panels B and D show a limited range of values. (C) Representative ECG waveform (average of 4 beats). (D) Waveform parameters are modulated following stimulus onset through the duration of heart rate excursion in the virally-injected mouse while parameters are unmodulated in non-injected controls. (Blue bar: 473nm, 5ms pulses, 20Hz, 10mW unmodulated power) (E) Quantification of ECG parameters during the pre-stimulus state (Pre), the period in which heart rate is perturbed > 1% from baseline (Stim), and the period following the heart rate perturbation (Post).

### Two-Photon Holographic Stimulation of Cervical Vagus Nerve in rAAV2-retro Heart-Injected Mice

Two-photon photostimulation of selected regions in the nerve was also performed. In addition to the heart-specific targeting achieved with retroviral delivery of the opsin, further selectivity is possible with spatial shaping of the excitation light. To this end, two-photon holographic photostimulation was employed. This approach enabled not only spatial selectivity of photoexcitation, but also reduced tissue scattering due to the longer wavelength excitation and thus greater access within the nerve^46^. The implantation of a GRIN lens-incorporated cuff on the cervical vagus nerve enabled coupling of the nerve to a movable-objective-microscope (MOM) for *in vivo* two-photon imaging and photostimulation. A spatial light modulator (SLM) integrated into the microscope beam path provided the capability to illuminate spatially selected regions of interest within the nerve^33^.

Region-specific photostimulation in the nerve yielded robust changes in cardiac function as well as respiration. An example of perturbation in breath rate along with breath distension is shown in Figure 6A. Differential responses in cardiac function were produced with differing spatial patterns of photostimulation. An example of this is presented in Figure 6C and D, in which a broad stimulation region spanning the nerve cross section resulted in a depressive heart rate response that led to temporary heart block (Figure 6C, asterisk), while more localized regions of stimulation caused an increase in heart rate along with an opposing set of ECG responses (Figure 6D).

**Figure 6:**
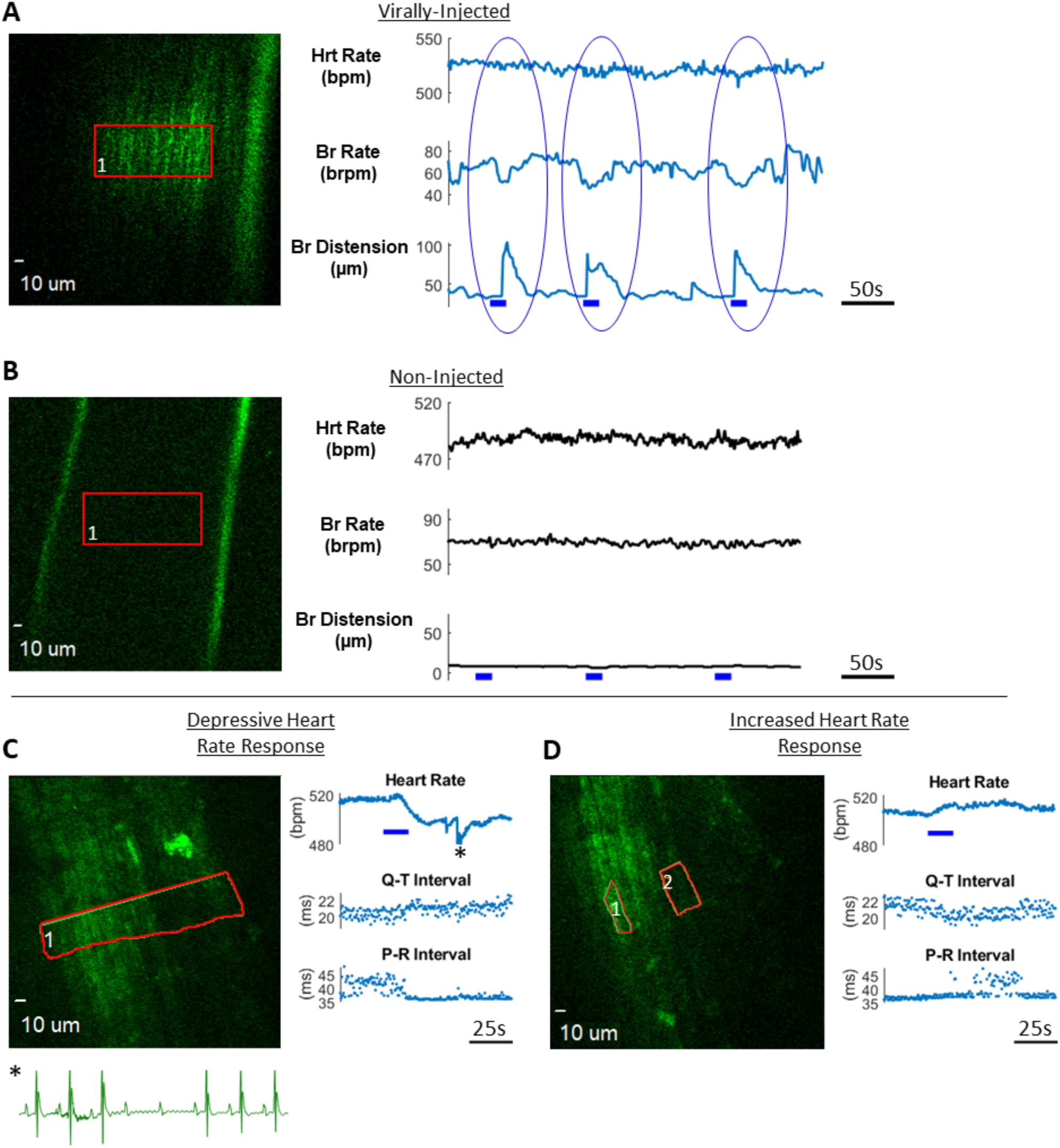
2-Photon holographic photostimulation in the cervical vagus nerve in mice infected with rAAV2-retro in the heart. (A) 920 nm photoexcitation (red ROI) in AAVrg-CAG-ChR2(h134R)-tdTomato-injected mouse elicited robust response in breath rate and distension (blue bar: 20Hz pulses, 5 ms pulse duration, 0.014 mW/μm^2^ unmodulated intensity). (B) Same photoexcitation in non-injected control mouse did not evoke any respiratory or cardiac responses. (C&D) 1030 nm photoexcitation in AAVrg-Syn-Chronos-eGFP-injected mouse elicited differential heart response with spatially distinct photoexcitation regions (red ROIs). The broad photoexcitation region in panel C caused a depressive heart rate response that led to heart block (asterisk, see inset) while the excitation profile in panel D caused an increase in heart rate and opposing ECG parameter changes. (blue bar: 20Hz pulses, 5 ms pulse duration, panel B: 0.12 mW/μm^2^ unmodulated intensity and panel C: 0.18 mW/μm^2^ unmodulated intensity).

## Discussion

The present study demonstrates highly improved targeting and specificity in peripheral nerve stimulation with optogenetic techniques. Holographic photostimulation was employed to stimulate subsets of vagal nerve fibers targeting the heart, resulting in drastically different cardiorespiratory responses depending on which fibers were stimulated (Figure 6). Furthermore, our results show the feasibility of targeted expression of opsins in vagal nerve fibers innervating the heart with a retrograde AAV virus injected into the myocardium. Cardiac viral injections elicited reporter expression in vagal axons, neurons within the nodose and superior cervical ganglia, and neuronal processes in the dorsal brainstem (Figures 3 and 4). Unexpectedly, we also found expression of the reporter in cholinergic neurons in NTS (Figure 4B). These findings demonstrate flexible alteration of cardiorespiratory function through targeted multiphoton cervical vagus holographic photostimulation. Additionally, the heart-specific opsin expression in vagal pathways achieved here with a retroviral approach will be applicable for neural targeting of other organ systems.

Vagal axons transduced by cardiac retrograde AAV delivery were determined to be primarily sensory afferents, since reporter expression was predominantly non-expressed in ChAT-driven fluorescently labeled fibers.. It should be noted that labeling was likely limited in part by regionally specific injection of the AAV. Along with axonal labeling we found neuronal expression in the nodose ganglion and abundant axonal expression in NTS. Labeling of sparse cholinergic somata was also found in NTS. This raises the question of whether expression in somata occurred due to direct axonal innervation to the heart from NTS neurons, or by trans-synaptic hopping of the rAAV2-retro from a sensory afferent of the nodose ganglion to the NTS neuron. To our knowledge there is no report of trans-synaptic hopping by rAAV2-retro (personal communication, Alla Karpova and David Schaffer^38^). Thus, it is possible although unlikely, that viral reporter in ChAT-positive neurons in NTS may reflect labeling of NTS neurons innervating the heart. If these neurons also make local synaptic connections, this would be consistent with the finding by Furuya and colleagues that acetylecholine injected into the cNTS increases phrenic frequency and affects sympathetic–respiratory coupling, without changing sympathetic activity^47^. Alternatively, uptake of the virus into the systemic or pulmonary circulation could possibly account for this unexpected expression. Future experiments are necessary to investigate this.

Photostimulation of the cervical vagus nerve resulted in changes in heart rate. Interestingly, both increases and decreases in heart rate were observed. This could be explained by differing afferent populations expressing the opsin across experiments, as well as spatially selective stimulation in the case of the two-photon holographic stimulation. While reduced heart rate (bradycardia) is often associated with vagal stimulation, afferent-dominated activation may lead to a suppression of efferent parasympathetic drive leading to increased heart rate^48,49^. ECG parameters were also perturbed during and following photostimulation. In addition to cardiac-specific indices, respiratory changes also occurred in response to stimulation, as measured with breath rate and distension. Brainstem cardiorespiratory modulatory circuits are well documented, and vagal afferents within NTS are known to drive respiratory centers, including the ventral respiratory column (VRC) within the medulla ^50,51^.

Two-photon holographic photostimulation was employed to selectively stimulate vagal activity. Using a custom microscope setup, and GRIN-lens integrated nerve cuff, this stimulation elicited the cardiac and respiratory effects described above, and also activated differential and opposing heart rate and ECG changes with distinct stimulation regions within the vagus nerve. This modality of photostimulation demonstrates an experimental tool which adds optical depth access in tissue, and selectivity for increased precision in studying the neural pathways affecting organ function.

Given that improved targeting is needed to precisely study and modulate organ function, we have investigated mechanisms within the optical (optogenetic) approach to neuromodulation. Delivery of opsins to organ-specific and genetically defined fibers is a critical aspect to this strategy. The present study provides an initial demonstration of retrograde opsin delivery in an organ-specific manner to facilitate peripheral nerve photomodulation. Results showed the feasibility of perturbing organ function with this technique. This work did not delve into the precise mechanisms and neuronal subsets giving rise to the cardiac and respiratory responses, however. Future studies may investigate defined subsets of peripheral autonomic fibers to determine and correlate neuronal molecular markers to distinct cardiac/organ functionalities. Identifying these neuronal markers of autonomic function and characterizing their effect on the organ will be central in the development of neuromodulatory strategies.

## Acknowledgements

We thank the following funding institutions that supported this work: National Institutes of Health, Stimulating Peripheral Activity to Relieve Conditions (SPARC) initiative NIH OT2OD023852 (RFW, JHC, EAG), NIH U01NS099577 (DR, EAG), National Science Foundation NSF CBET-1631912 (EAG, DR), and Rocky Mountain Regional VA Medical Center R&D Research Career Scientist Award IK6RX002996 (RFW).

## Author Contributions

AKF performed the vagal stimulation experiments, tissue processing & imaging, analyzed the experimental data and drafted the manuscript. GLF built, aligned and characterized the 2-photon microscope and holography systems and operated these systems during experiments. PSR performed the cardiac viral-injection surgeries and provided input on the manuscript. SL fabricated the GRIN-cuff device and participated in experiments. NM processed the brain tissue and performed immunohistochemical staining of tissues. KS and JLA provided support and cardiac expertise in the investigation. DR and JHC advised on viral constructs and transgenic animals. EAG oversaw development of the 2-photon microscopy and holography aspects of the project and assisted with drafting the Methods section. RFW oversaw the development and fabrication of the nerve cuff device. RFW, EAG, JHC, and DR initiated and directed the study and edited the manuscript.

