## Supplementary Figures for "Selective Modulation of Heart and Respiration by Optical Control of Vagus Nerve Axons Innervating the Heart"

**Supplemental Figures**

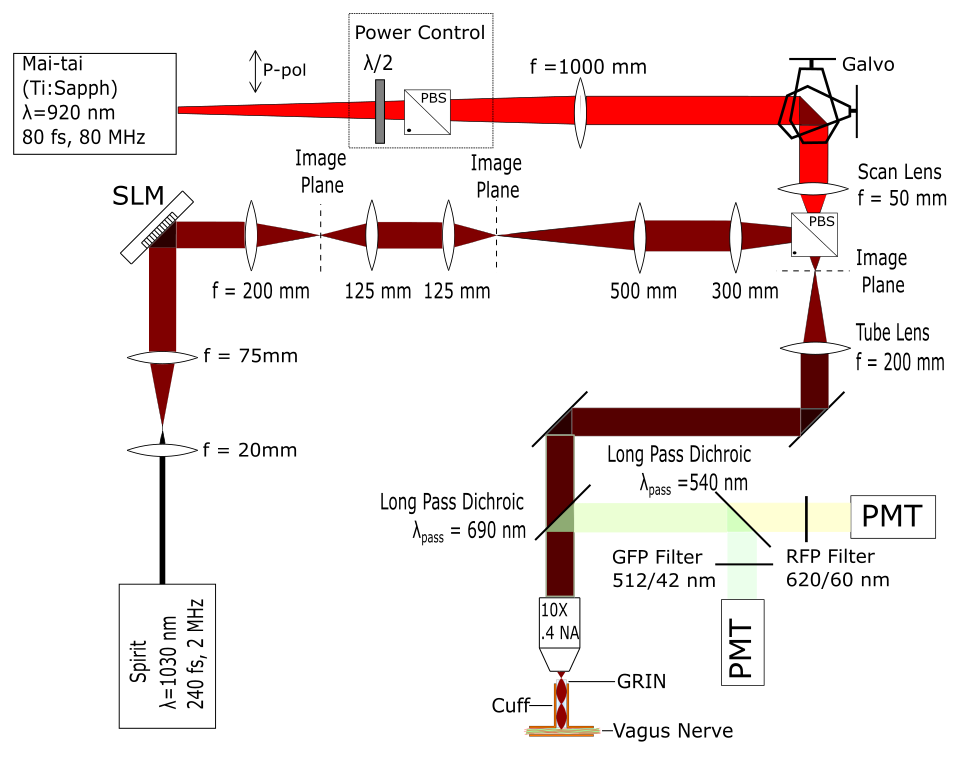

Supplementary Figure 1: Diagram of two-photon imaging and holographic photostimulation setup showing the beam path for imaging and a separate path for photostimulation using a spatial light modulator for shaping the intensity pattern at the focus. The beams are combined after the scan lens using a polarizing beam splitter. The GRIN-cuff is positioned under the air objective to focus into the cervical vagus nerve.

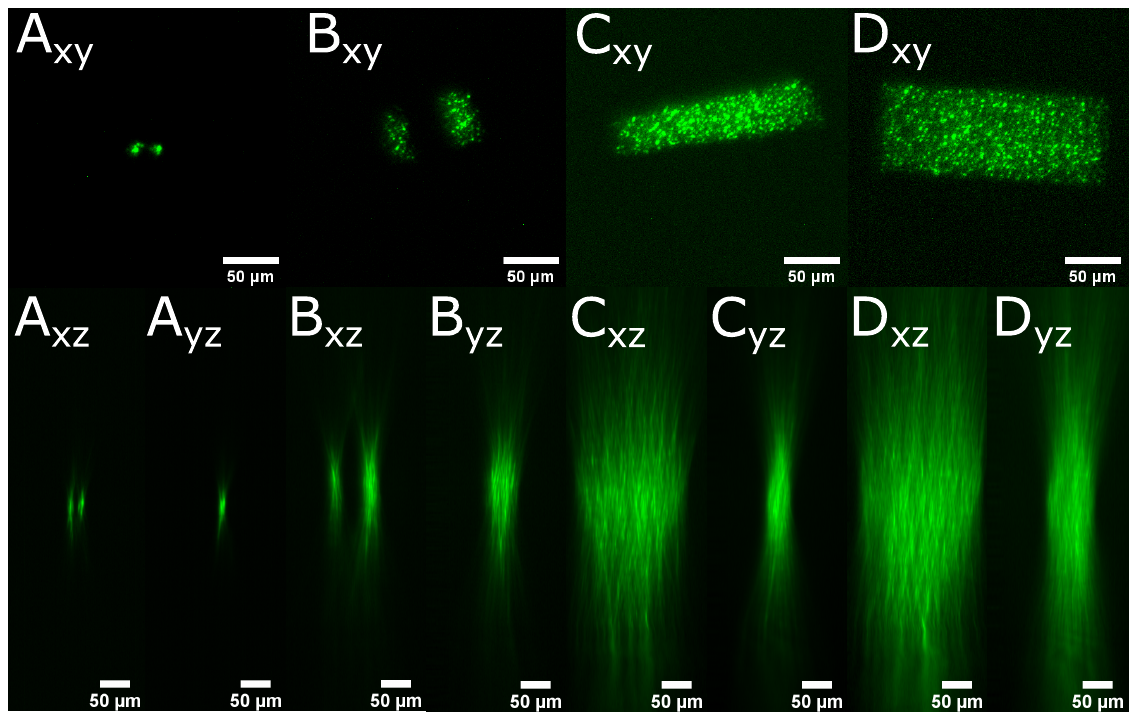

Supplementary Figure 2: Transverse (xy) and axial (xz, yz) profiling of holographic two-photon excitation at the focus of the GRIN relay lens within the vagus nerve. (A) spatial profile for two 10 μm spots (B-D) excitation profiles used for the *in vivo* studies reported in Fig. 6. The axial FWHM were calculated from the z stacks to be (A) 44 microns, (B) 105 microns (left region) 140 microns (right region), (C) 179 microns, and (D) 344 microns. Images were acquired by translating the two-photon holograms being formed at the object plane of the GRIN lens through a thin fluorescent slide with emission imaged with stationary inspection microscope onto a camera.

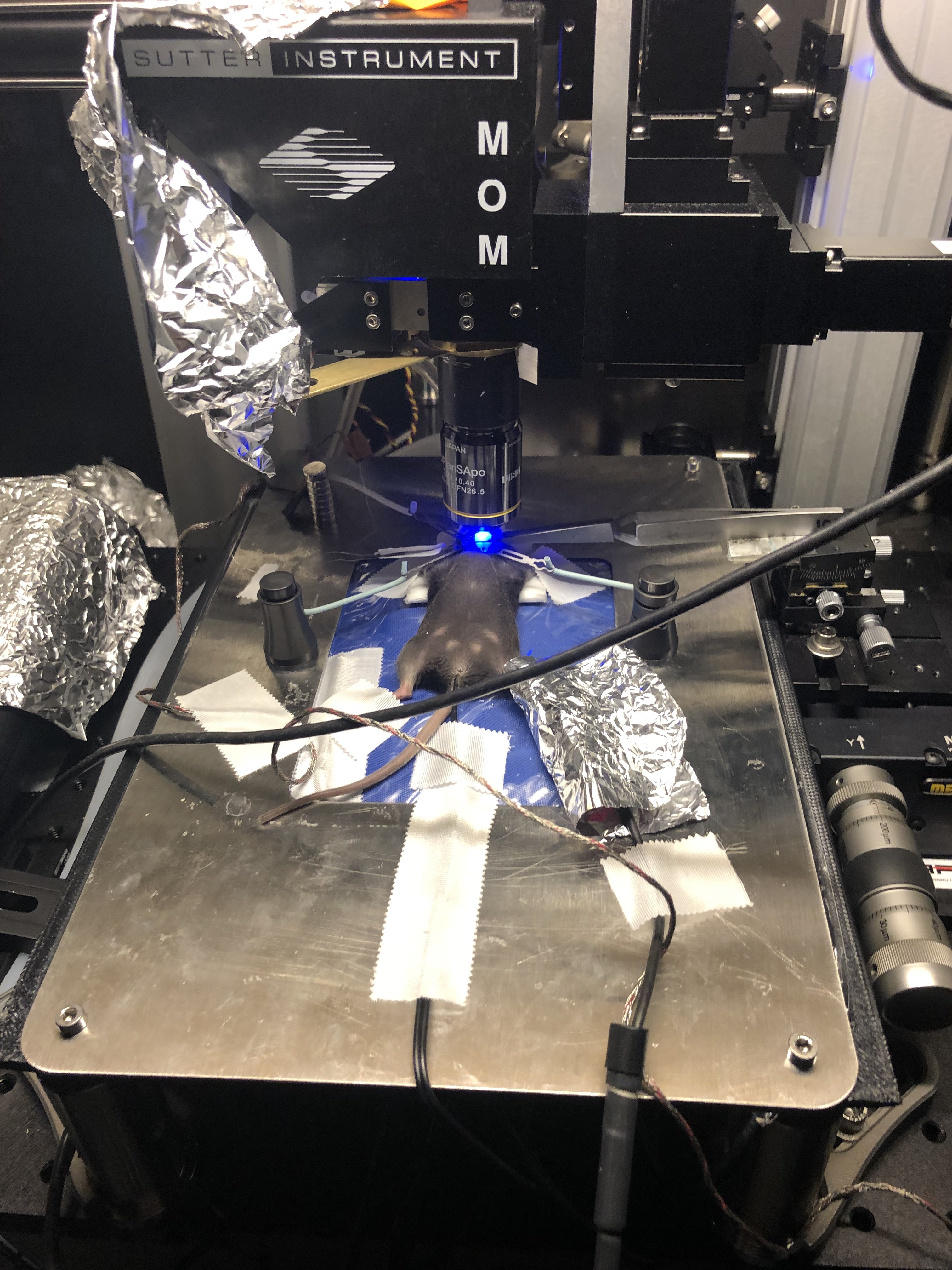

Supplementary Figure 3: Experimental setup of mouse two-photon imaging and photostimulation. Anesthetized mouse is supine on a heating pad with a pulse oximetry paw sensor for vitals measurement. The GRIN lens is held upright using custom forceps on a mechanical stage. The objective is positioned above the GRIN lens for optical access of the vagus nerve.
